# Hierarchical Flows of Human Cortical Activity

**DOI:** 10.64898/2026.03.19.712872

**Authors:** Xiaobo Liu, Alex I. Wiesman, Sylvain Baillet

## Abstract

Ongoing brain activity unfolds as structured spatiotemporal patterns across the cortex, yet quantifying the direction and strength of this propagation on the folded cortical sheet is challenging within and across individuals. We introduce geodesic *cortical flow*, a surface-based optical-flow framework that estimates millisecond-resolved surface-tangent propagation fields from source-imaged magnetoencephalography (MEG) data. In resting-state MEG from 608 healthy adults, spontaneous propagation was anisotropic and bidirectionally aligned with the principal unimodal-to-transmodal functional gradient: slow activity (1-13 Hz) was biased toward upstream propagation from sensory to association cortex, whereas beta activity (13-30 Hz) was biased toward downstream propagation in the opposite direction. Across adulthood, this balance shifted toward weaker upstream slow propagation and stronger downstream beta propagation. Propagation strength, indexed by kinetic energy of the cortical flow, followed a robust posterior-to-anterior gradient and, within frontoparietal cortex, higher kinetic energy was associated with better fluid intelligence after adjustment for age. Kinetic-energy dynamics further identified stable-state dwell times that tracked regional neuronal timescales. Together, these findings establish geodesic cortical flow as a geometry-informed framework for quantifying frequency-resolved cortical propagation and its variation across aging and cognition.

## Introduction

Spontaneous brain activity unfolds as coordinated spatiotemporal patterns that reflect the interaction of local circuit dynamics and long-range connectivity (Makeig et al., 2009; Roberts et al., 2019; Sporns et al., 2005). Invasive recordings have shown that cortical activity can propagate as traveling waves across the folded surface during perception, sleep, and pathological states (Massimini et al., 2004; Muller et al., 2014, 2018; Smith et al., 2022). Complementary fMRI work has identified slower whole-cortex propagation motifs, including bidirectional patterns aligned with the principal functional gradient from unimodal sensory regions to transmodal association cortex (Margulies et al., 2016; Pines et al., 2023). Whether comparable hierarchy-aligned propagation is expressed at the millisecond timescale of human neurophysiological activity remains unresolved.

Here we address this question using source-imaged magnetoencephalography (MEG) and a surface-based optical-flow framework adapted to the cortical manifold (Baillet, 2017; Lefèvre & Baillet, 2008, 2009). This approach estimates local propagation vectors directly on the folded cortical sheet, allowing direction and magnitude to be quantified in anatomically meaningful coordinates. We focus on slow (1–13 Hz) and beta (13–30 Hz) activity because large-scale cortical dynamics vary across frequency, and prior work links slower and beta-band activity to partially distinct modes of sensory, integrative, and control-related processing (Engel et al., 2013; Klimesch, 2018; Lundqvist et al., 2018; Richter et al., 2017; Siclari et al., 2017).

We further tested whether these propagation motifs are reorganized across the adult lifespan. Based on evidence for age-related changes in sensory reliability and top-down control (Engel-Yeger et al., 2021; Fontolan et al., 2014; Jones & Noppeney, 2021; Son et al., 2023; Yang et al., 2023), we predicted a frequency-specific shift, with weaker upstream propagation in slow activity and stronger downstream propagation in beta activity in older adults.

## Results

### Cortical flow framework

We modeled millisecond-resolved MEG source activity as a time series of scalar maps on the cortical mesh and estimated surface-tangent propagation vectors using an optical-flow method adapted to curved manifolds (Lefèvre & Baillet, 2008, 2009). At each vertex and time point, the method returns a tangent-plane vector whose orientation reflects the direction of maximal local spatiotemporal change and whose magnitude reflects instantaneous propagation strength (Fig. 1A).

**Figure 1:**
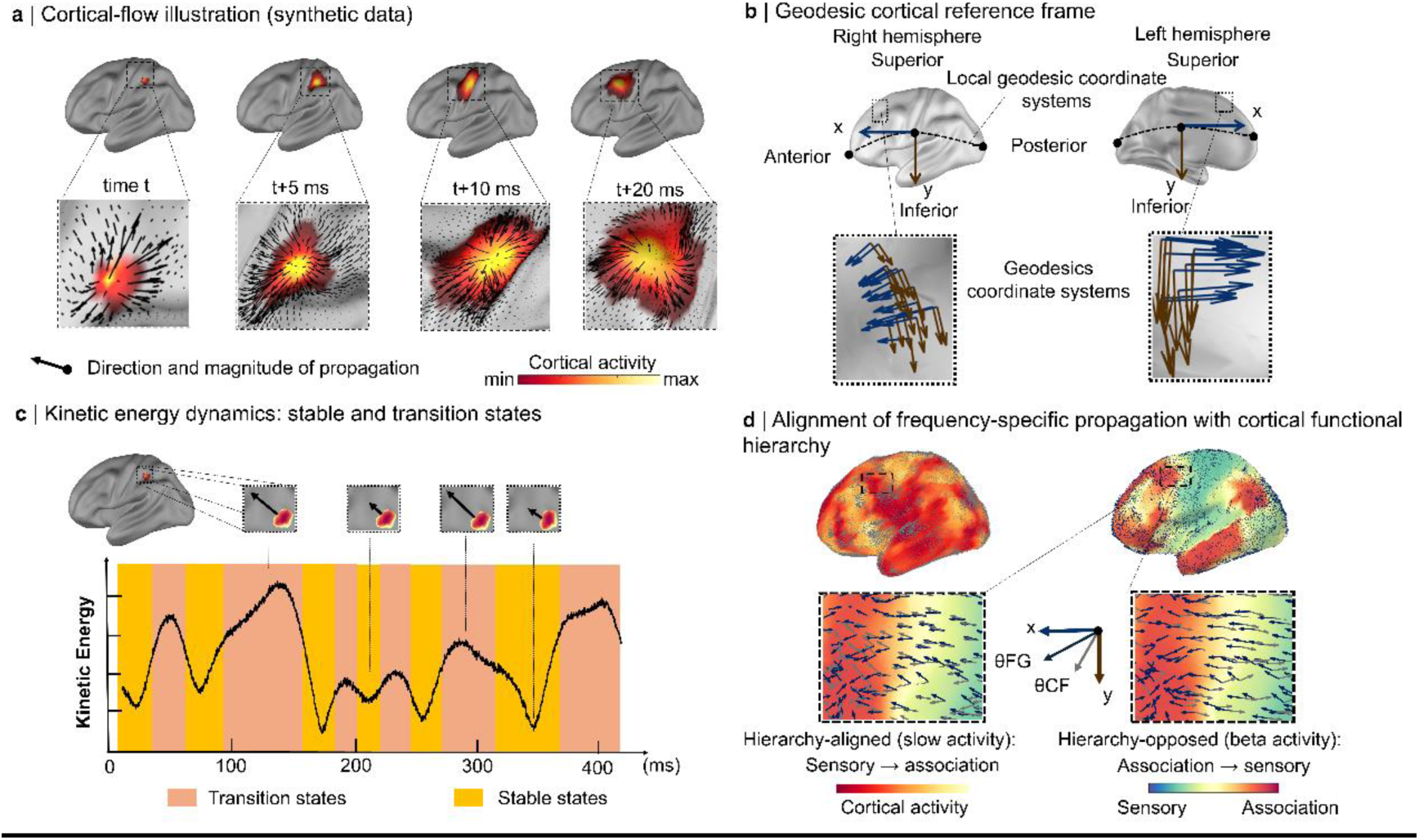
Geodesic cortical-flow framework and kinetic-energy states. **a | Cortical-flow:** Surface-based optical flow estimates a surface-tangent propagation vector at each cortical vertex and time point from source-imaged MEG activity. Black arrows indicate the local direction and magnitude of propagation (example frames from synthetic data). **b | Geodesic reference frame:** For each hemisphere and at each cortical location, a local tangent plane defines orthogonal geodesic axes aligned with canonical anatomical directions. Blue arrows indicate the posterior→anterior axis and brown arrows the superior→inferior axis, enabling consistent measurement of propagation direction across the folded cortical surface. **c | Kinetic energy dynamics**: Kinetic energy, defined as the squared magnitude of the cortical-flow vector, fluctuates over time between low-energy stable states and high-energy transition states. Example epochs from synthetic data and the corresponding global kinetic-energy trace are shown. **d | Alignment of frequency-specific propagation with cortical functional hierarchy:** The local direction of the principal functional gradient defines θ_FH_, and the cortical-flow direction defines θ_CF_. The unsigned angular difference Δθ between θ_CF_ and θ_FH_ classifies propagation as hierarchy-aligned or hierarchy-opposed, corresponding preferentially to slow and beta activity, respectively.

To interpret directions on the folded surface, we expressed these vectors in a local geodesic reference frame defined on each vertex’s tangent plane, with orthogonal posterior-anterior and superior-inferior axes (Fig. 1B; Le Troter et al., 2012). All reported angles are therefore referenced to an anatomically grounded coordinate system rather than to Euclidean directions in volume space.

We summarized propagation strength using kinetic energy, defined as the squared magnitude of the cortical-flow vector (Methods). Over time, kinetic energy alternated between low-energy epochs characterized by relatively coherent, slowly varying propagation and higher-energy epochs marked by rapid changes in vector direction and magnitude (Fig. 1C).

To quantify alignment with cortical hierarchy, we compared the direction of cortical flow at each vertex with the local direction of the principal unimodal-to-transmodal functional gradient (Margulies et al., 2016). We classified flow as *upstream* when it aligned with the sensory-to-association axis and *downstream* when it opposed that axis (Fig. 1D).

We assessed the geometric validity of this framework in two complementary ways. Synthetic simulations with known propagation trajectories showed that cortical flow accurately recovered canonical anatomical directions, including trajectories that followed curved cortical paths (Supplementary Fig. 3). In parallel, the anatomy-informed geodesic reference frame yielded consistent directional interpretation across the folded cortical surface and across participants (Supplementary Fig. 2), indicating that the directional effects reported below are unlikely to be artifacts of cortical geometry or coordinate choice.

### Spontaneous activity propagates along the cortical functional hierarchy

Across resting-state recordings from 608 adults, broadband cortical flow exhibited a dominant posterior-to-anterior orientation (Fig. 2A). Here, broadband refers to 0.6 Hz up to each participant’s 95% PSD cut-off frequency (mean ≈ 80 Hz; Methods). The group-mean flow angle was 72.22° (95% CI, 71.40°–73.00°), and participant-level mean directions departed significantly from uniformity (Hodges-Ajne test, p < 0.05). To ensure that this result was not produced by averaging across opposing directions, we verified the same bias with an incidence-based anatomical-axis metric contrasting posterior-to-anterior and anterior-to-posterior events at the participant level (Supplementary Fig. 1).

**Figure 2.**
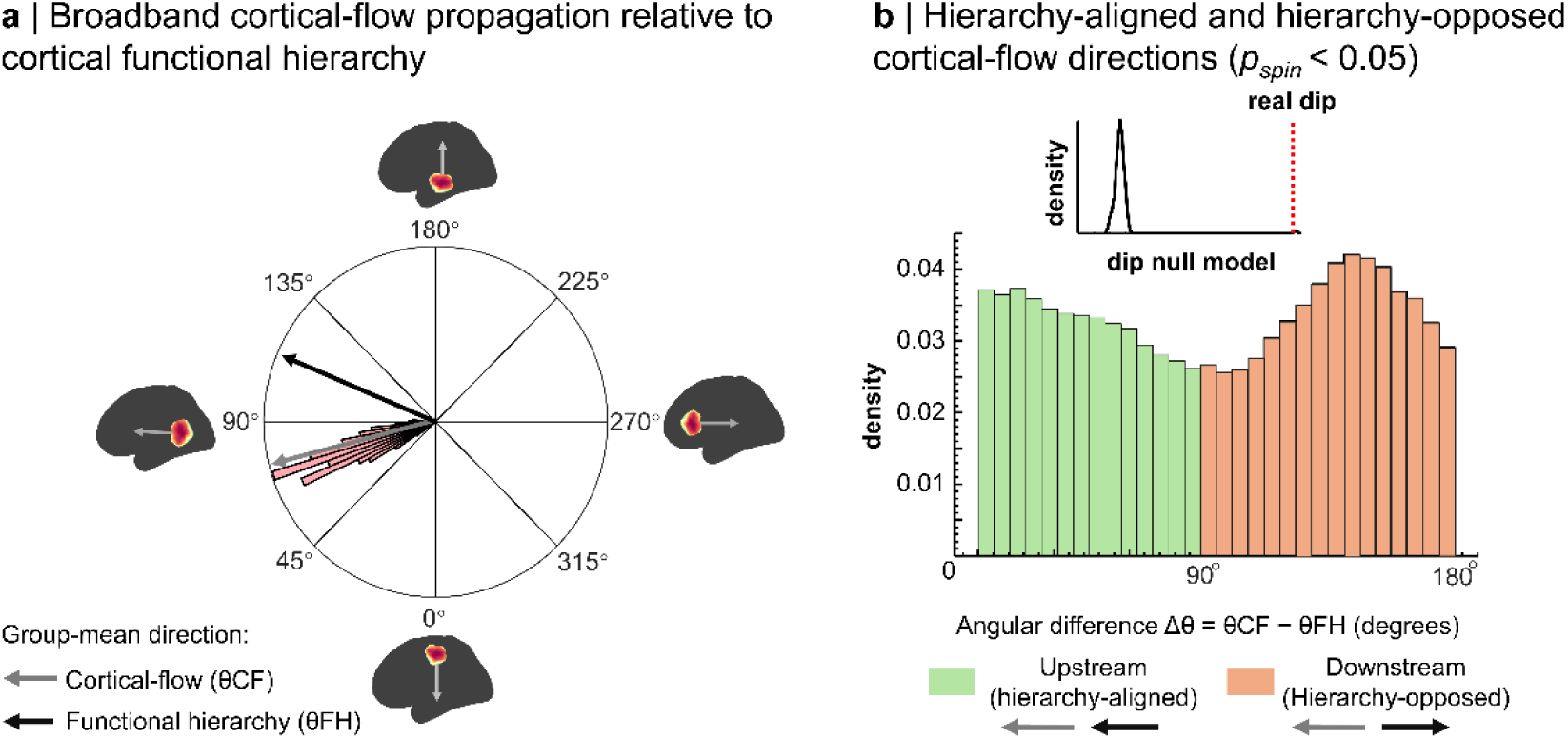
Spontaneous cortical flow aligns with the cortical functional hierarchy. **a | Dominant direction of broadband cortical flow relative to the cortical functional hierarchy:** The rose plot shows each participant’s mean cortical-flow direction (θ_CF_). The group-mean direction (gray arrow; 72°) closely parallels the mean orientation of the principal functional hierarchy (θ_FH_; black arrow). Insets illustrate representative cortical-flow and hierarchy directions projected onto inflated cortical surfaces. **b | Angular relationship between cortical-flow direction and the functional hierarchy:** Histogram shows vertex-wise angular differences between cortical-flow direction and hierarchy orientation (Δθ = θ_CF_ − θ_FH_), pooled across participants and time points. Angles near 0° (green) indicate hierarchy-aligned propagation, whereas angles near 180° (orange) indicate hierarchy-opposed propagation. Inset shows the empirical Hartigan dip statistic (red dashed line) relative to a null distribution generated from 1000 spin permutations, confirming a non-uniform, bimodal alignment (*p_spin_* < 0.05).

We next asked how these directions were oriented relative to cortical functional hierarchy. The angular difference between cortical-flow direction and the local hierarchy axis (Δθ = θCF − θFH) was strongly bimodal, with one mode near 0° (upstream; sensory-to-association) and the other near 180° (downstream; association-to-sensory) (Fig. 2B). Hartigan’s dip test rejected unimodality relative to spin-permutation nulls both for participant-averaged angles and for the full set of time-point angles (both pspin < 0.001), indicating robust bidirectional propagation aligned with the principal functional gradient.

### Frequency-specific and age-dependent directionality

Band-limited analyses revealed a marked frequency dependence of hierarchy alignment (Fig. 3A). Across participants, upstream propagation was more prevalent in slow activity than in beta activity (paired t(607) = 17.9, p < 0.001, Cohen’s d = 0.73), demonstrating that the balance between upstream and downstream propagation depends strongly on spectral content.

**Figure 3.**
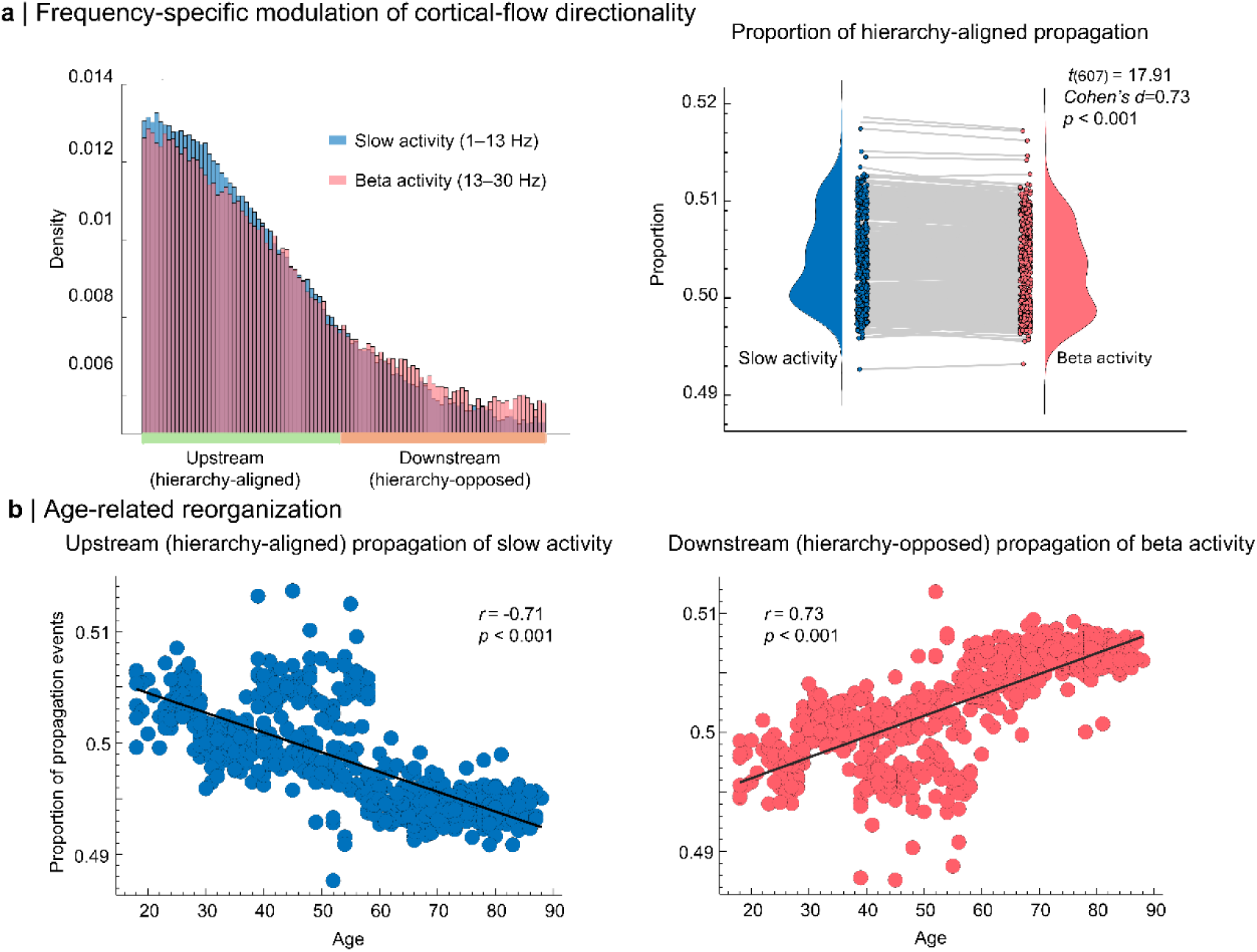
Frequency- and age-dependent modulation of cortical-flow directionality. **a | Frequency-specific directionality of cortical flow relative to the functional hierarchy:** Left: histogram of vertex-wise angular differences between cortical-flow vectors and the functional hierarchy (Δθ = θ_CF_ − θ_FH_), pooled across participants and time points. Slow activity (1–13 Hz; blue) exhibits a greater incidence of hierarchy-aligned propagation, whereas beta activity (13–30 Hz; red) is biased toward hierarchy-opposed propagation. Right: paired violin and box plots show, for each participant, the proportion of hierarchy-aligned propagation events. This proportion is significantly higher for slow than for beta activity (*t*(607) = 17.91, *p* < 0.001, *Cohen’s d* = 0.73). **b | Age-related reorganization of frequency-specific propagation:** The proportion of hierarchy-aligned propagation in slow activity decreases with age (left; *r* = −0.71, *p* < 0.001), whereas the proportion of hierarchy-opposed propagation in beta activity increases with age (right; *r* = 0.73, *p* < 0.001). Together, these results demonstrate a frequency-specific propagation bias relative to cortical functional hierarchy that systematically reorganizes across the adult lifespan.

Propagation direction also changed systematically with age (Fig. 3B). The proportion of upstream propagation in slow activity declined with age (r = −0.71, p < 0.0001), whereas the proportion of downstream propagation in beta activity increased with age (r = 0.73, p < 0.0001). These effects remained significant in covariate-adjusted models that included sex, handedness, head motion, and residual ocular and cardiac components (all pFDR < 0.05; Supplementary Fig. 1), indicating that lifespan effects were robust to these nuisance factors.

### Kinetic-energy hierarchy and its reorganization with age

Propagation strength exhibited a clear hierarchical organization. Vertex-wise broadband kinetic energy was highest in posterior sensory cortex and lowest in association cortex, yielding a strong negative spatial association with functional-hierarchy rank (r = −0.66, pspin < 0.001; Fig. 4A). Within frontoparietal cortex, higher kinetic energy was positively associated with fluid intelligence after adjustment for age (r = 0.28, p < 0.001; Fig. 4B), linking propagation strength in association networks to inter-individual differences in cognitive performance.

**Figure 4.**
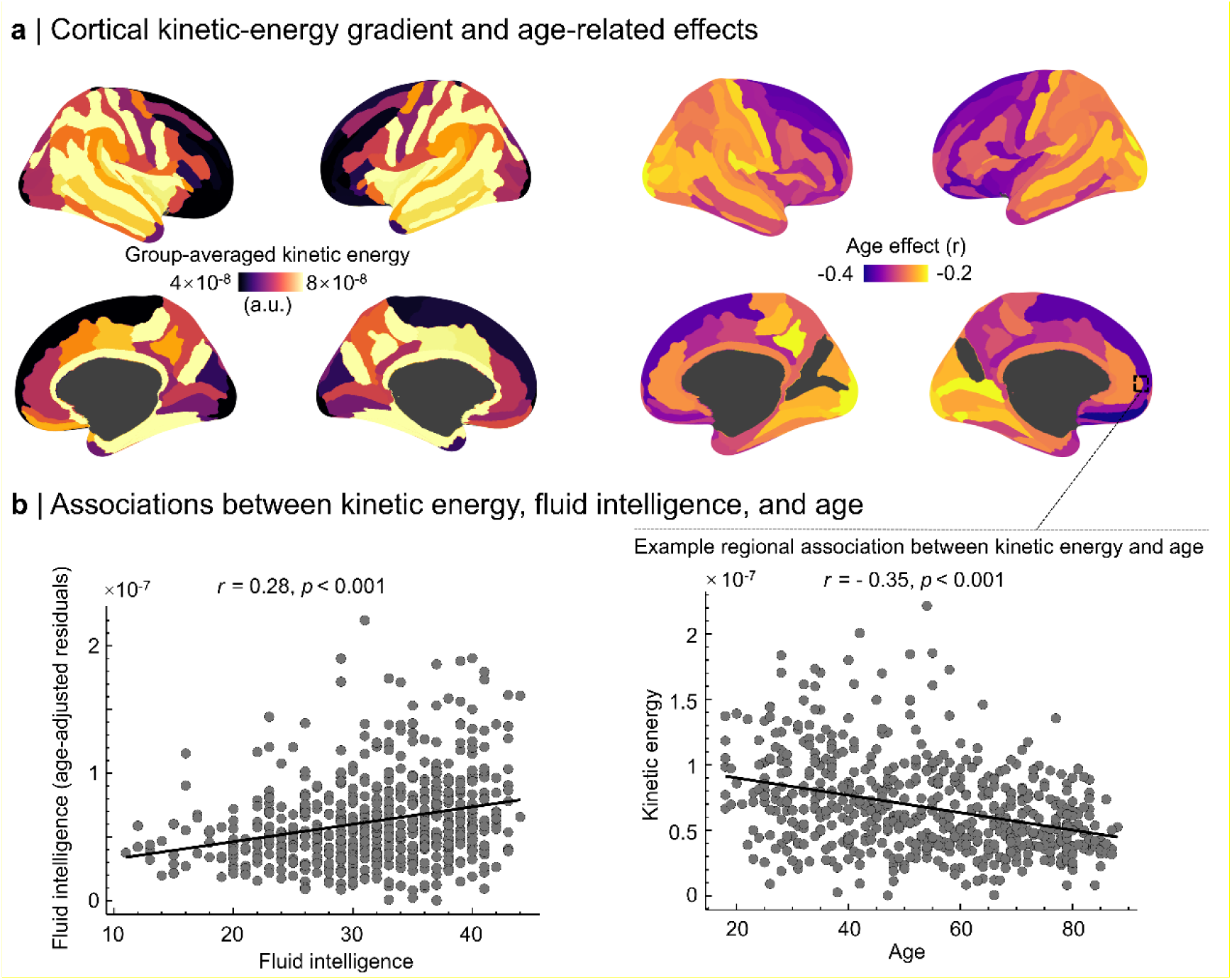
Cortical kinetic energy tracks functional hierarchy, aging, and cognition. **a | Cortical kinetic-energy gradient and age-related effects:** Left: group-averaged kinetic-energy maps reveal a posterior→anterior decrease in propagation strength (4 × 10⁻⁸ to 8 × 10⁻⁸ a.u.). Right: vertex-wise correlations with age show predominantly negative effects in frontal cortex. Inset highlights the relationship between frontoparietal kinetic energy and age (*r* = −0.35, *p* < 0.001). **b | Associations between cortical kinetic energy, fluid intelligence, and age**: Left: frontoparietal kinetic energy correlates positively with fluid intelligence after controlling for age (*r* = 0.28, *p* < 0.001). Right: example regional association illustrating the decline of kinetic energy with age in orbitofrontal cortex (same data as inset in panel a). Each dot represents one participant.

Kinetic-energy maps remained structured in both the slow and beta bands (Fig. 5A). The beta-to-slow kinetic-energy ratio, used here as a frequency-balance index, was higher in association cortex than in sensory cortex and varied systematically along the functional hierarchy (r = −0.66, pspin < 0.05). Aging produced a frequency-specific reorganization of this balance (Fig. 5B): global kinetic energy declined with age (pFDR < 0.05), with the strongest reductions in orbitofrontal and frontoparietal regions (peak r = −0.35), whereas the beta-to-slow ratio decreased with age in association cortex and increased in posterior regions (peak r = 0.39, pFDR < 0.001). The spatial pattern of age effects also tracked hierarchy rank (r = −0.49, pspin < 0.05), indicating graded kinetic reorganization along the sensory-to-association axis.

**Figure 5.**
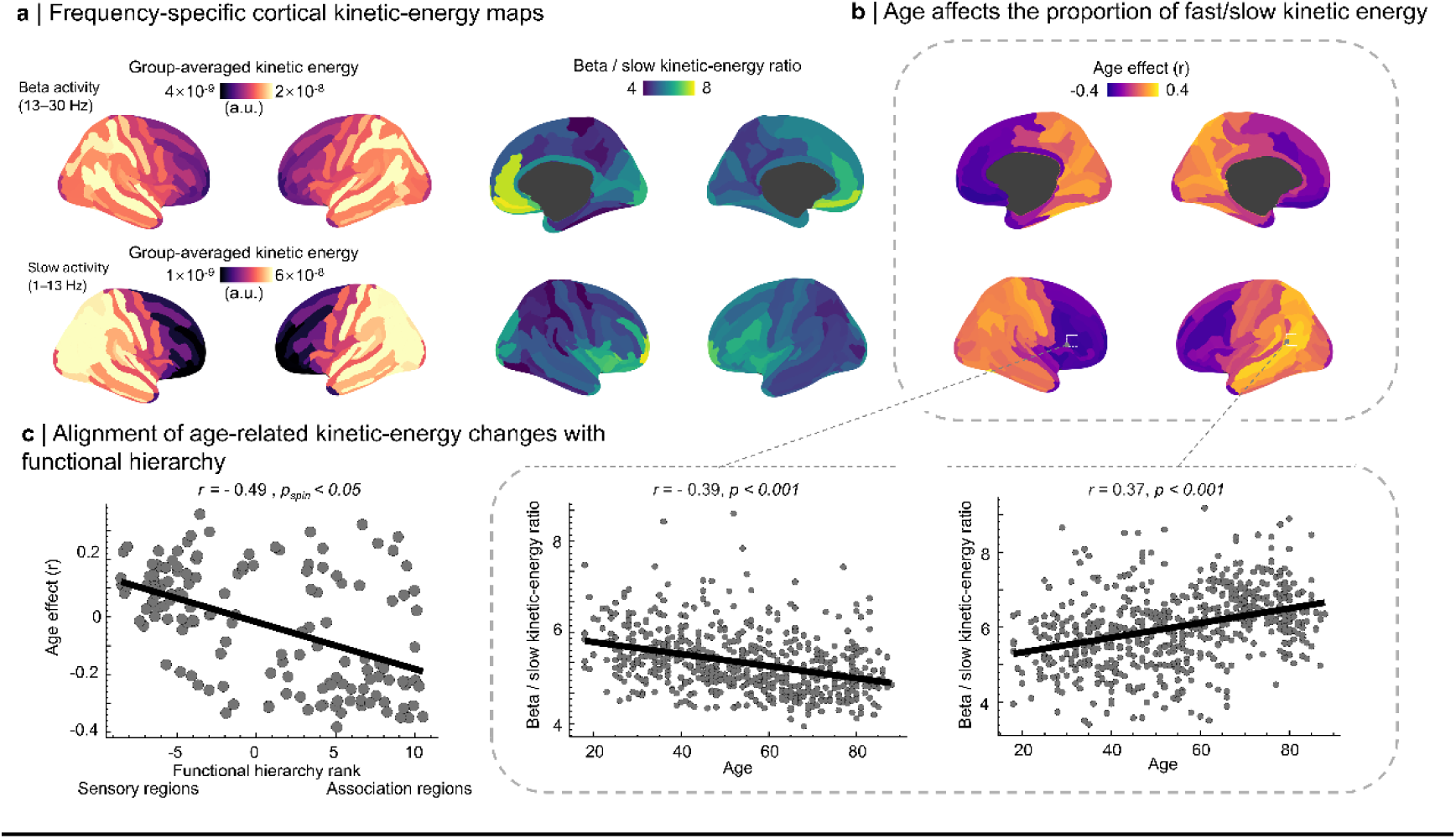
Frequency-specific topography of cortical kinetic energy: age dependence and alignment with functional hierarchy. **a | Frequency-specific cortical kinetic-energy maps:** Top row: beta activity (13–30 Hz). Bottom row: slow activity (1–13 Hz). Both frequency bands exhibit a posterior→anterior gradient in kinetic energy. Center panels show the beta-to-slow kinetic-energy ratio, which is highest in anterior cortex and inversely related to functional hierarchy rank (*r* = −0.66, *p_spin_* < 0.05). **b | Age-related modulation of the beta-to-slow kinetic-energy ratio**: Vertex-wise correlations reveal opposing age effects across the cortex, with relative decreases in anterior regions and increases in posterior regions (*p_FDR_* < 0.05). Insets illustrate representative anterior (inferior frontal gyrus; *r* = −0.39, *p* < 0.001) and posterior (inferior parietal gyrus; *r* = 0.37, *p* < 0.001) regions. **c | Alignment of age-related kinetic-energy changes with functional hierarchy:** Vertex-wise age coefficients are negatively correlated with functional hierarchy rank (*r* = −0.49, *p_spin_* < 0.05), indicating that age-related changes in kinetic energy balance systematically follow the sensory→association hierarchy. Each dot represents one *Destrieux* parcel.

### Stable and transient kinetic states

Kinetic-energy time series enabled a complementary time-domain characterization of cortical stability by separating *stable* states from *transition* states. Stable states were defined around local minima of kinetic energy and transition states around local maxima after band-specific smoothing (Methods). Stable-state dwell times increased modestly but systematically from posterior unimodal to anterior association cortex (Fig. 6A): dwell times were longer in association regions such as orbitofrontal cortex (81 ± 21 ms) and lateral prefrontal cortex (79 ± 21 ms) than in unimodal regions such as primary auditory cortex (75 ± 19 ms), primary somatosensory cortex (77 ± 20 ms), and MT (77 ± 20 ms). Across parcels, stable-state dwell time correlated with intrinsic neuronal timescales estimated from spectral knee parameters (r = 0.49, pspin < 0.0001; Fig. 6B) and increased with cortical hierarchy rank (pspin < 0.05), linking time-domain stability to established cortical temporal hierarchies.

**Figure 6.**
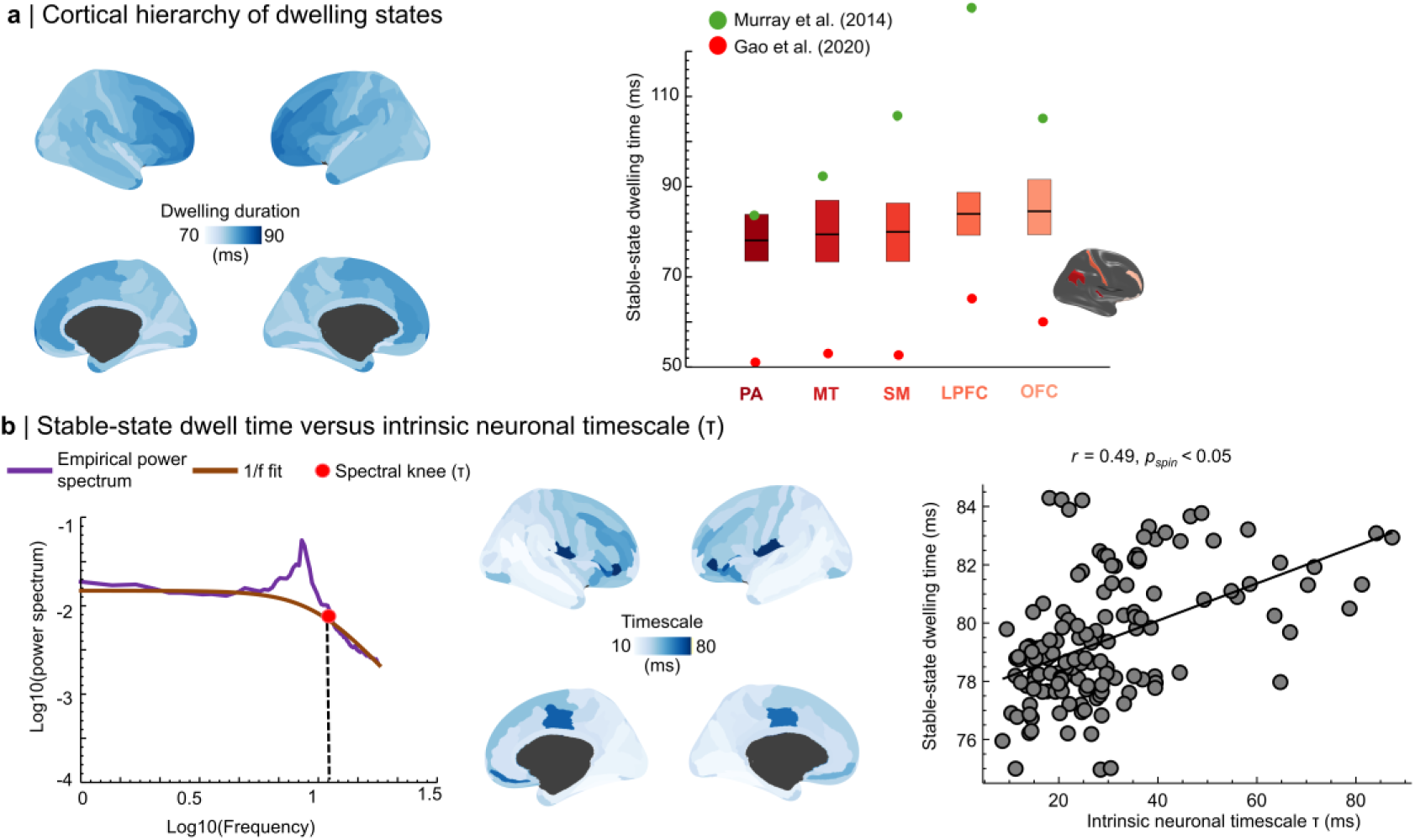
Stable-state dwell times align with cortical temporal hierarchy and intrinsic neuronal timescales. **a | Cortical hierarchy of stable-state dwell time:** Surface maps show median stable-state dwell times derived from local kinetic-energy fluctuations, revealing a posterior→anterior increase across the cortex. Bar plots summarize representative parcels (PA, primary auditory; MT, middle temporal; SM, primary somatomotor; LPFC, lateral prefrontal cortex; OFC, orbitofrontal cortex), demonstrating longer dwell times in association cortex. These spatial patterns are consistent with previously reported cortical temporal hierarchies (Murray et al., 2014; Gao et al., 2020). **b | Stable-state dwell time versus intrinsic neuronal timescale (τ):** Left: example power spectrum illustrating the 1/f fit and spectral knee defining τ. Center: cortical map of τ values. Right: scatter plot shows a positive association between stable-state dwell time and intrinsic neuronal timescale across cortical regions (*r* = 0.49, *p_spin_* < 0.0001).

Stable-state dwell times also showed frequency-specific organization. Across the cortex, slow activity exhibited longer stable periods than beta activity, indicating greater temporal persistence of slow-frequency propagation patterns. The beta-to-slow dwell-time ratio showed a structured spatial distribution and systematic age dependence (pFDR < 0.05; Supplementary Fig. 6), suggesting that fast and slow propagation dynamics are rebalanced across adulthood.

### Replication in an independent dataset

To assess generalizability, we repeated the core analyses in an independent resting-state cohort from the OMEGA dataset (N = 83; Niso et al. 2016). Despite differences in acquisition parameters and sample composition, broadband cortical flow again showed a dominant posterior-to-anterior orientation that closely paralleled the principal functional hierarchy. Angular differences between cortical-flow direction and the hierarchy were again strongly bimodal (pspin < 0.001), and slow activity again showed a higher prevalence of hierarchy-aligned propagation than beta activity. These replication results indicate that the frequency-dependent directional structure of cortical flow is reproducible across datasets and MEG acquisition systems (Supplementary Figs. 4–6).

## Discussion

Using a geodesic cortical-flow framework applied to source-imaged MEG, we show that spontaneous cortical propagation is structured, frequency-dependent, and aligned with the brain’s macroscale functional hierarchy. Resting-state activity did not propagate isotropically across the cortical surface. Instead, propagation was bidirectionally organized along the principal unimodal-to-transmodal gradient, with slow activity biased toward upstream propagation and beta activity biased toward downstream propagation. Across adulthood, this balance shifted toward weaker upstream slow propagation and stronger downstream beta propagation. In parallel, kinetic energy and stable-state dwell times revealed complementary gradients in propagation strength and temporal stability. Together, these findings establish cortical flow as a compact, geometry-aware, frequency-resolved description of spontaneous human cortical dynamics.

### Cortical propagation aligns with anatomical and functional hierarchy

At rest, spontaneous propagation preferentially followed reciprocal posterior-to-anterior and anterior-to-posterior axes, consistent with structured bidirectional interactions between sensory and association cortex. This extends reports of hierarchy-aligned propagation from intracranial recordings and time-resolved fMRI to millisecond-scale human electrophysiology (Pines et al., 2023; Zhang et al., 2018). The replication of these effects across independent datasets and MEG systems argues against a trivial explanation based on coordinate choice, averaging artifacts, or a single preprocessing pipeline.

A likely substrate is that multiple structural features vary systematically along the sensory-to-association axis, including gradients in myeloarchitecture, laminar differentiation, and neuronal density (Cahalane et al., 2012; Charvet et al., 2014, 2015; Huntenburg et al., 2018; Valk et al., 2020). These gradients do not by themselves determine propagation direction, but they provide a biologically plausible anatomical context in which hierarchy-aligned propagation can emerge and remain spatially organized.

### Frequency-specific directionality suggests complementary hierarchical modes

The dissociation between upstream slow activity and downstream beta activity suggests complementary propagation modes with distinct spectral content. This pattern is consistent with theories that link slower activity to broad coordination across cortical systems and beta-band activity to feedback-related or context-setting dynamics (McCormick & Bal, 1994; Shine, 2019; Shine, 2021; Ujma et al., 2022; Baillet, 2017; Dubey et al., 2023). At the same time, our measures quantify the propagation of band-limited activity patterns rather than directed synaptic communication per se. We therefore interpret these effects as frequency-specific dynamical motifs that are compatible with, but do not by themselves prove, bottom-up and top-down signaling accounts.

### Aging rebalances frequency-specific propagation along the hierarchy

Across adulthood, these frequency-specific modes were systematically reweighted: upstream slow propagation weakened with age, whereas downstream beta propagation strengthened. This pattern is consistent with prior evidence that aging is accompanied by reduced reliability of sensory representations and greater reliance on internally generated or top-down influences (Fernandez-Ruiz et al., 2018; Jones & Noppeney, 2021; Toussaint et al., 2014). Because the present data are cross-sectional, however, these age effects should not be overinterpreted as developmental trajectories in a strict causal sense; compensation, dedifferentiation, and cohort effects remain plausible contributors.

### Kinetic energy and temporal stability provide complementary markers

Beyond directional organization, kinetic energy revealed a robust posterior-to-anterior gradient in propagation strength, with the highest values in posterior sensory cortex and the lowest values in transmodal association cortex. Supplementary analyses further suggested an association between kinetic energy and cortical myelination, consistent with sensitivity to known microstructural gradients. In contrast, temporal stability was better captured by stable-state dwell times, which increased toward association cortex and tracked intrinsic neuronal timescales estimated from spectral knees. Taken together, these findings suggest that propagation strength and propagation stability capture complementary aspects of cortical organization rather than a single common gradient.

One possible interpretation is that higher kinetic energy in sensory cortex reflects greater responsiveness to rapidly changing activity patterns, whereas longer dwell times in association cortex reflect more persistent, integrative dynamics. We treat this interpretation cautiously: kinetic energy and dwell time are descriptive summaries of propagation fields, not direct measures of synaptic gain or circuit mechanism. Even so, their orderly spatial structure and behavioral associations indicate that cortical flow captures biologically meaningful variation rather than only phenomenological texture.

### Limitations

Several limitations should be noted. First, hierarchy alignment was defined relative to a normative fMRI-derived functional gradient (Margulies et al., 2016), which does not capture subject-specific or age-dependent differences in hierarchical organization; deriving participant-specific gradients is an important next step. Second, although the geodesic framework explicitly respects cortical geometry, other anatomical features such as myelin, curvature, sulcal depth, and cortical thickness likely influence propagation trajectories and should be tested directly. Third, the mechanisms underlying the observed directionality—local circuit dynamics, long-range axonal coupling, subcortical inputs, and geometric constraints—cannot be disentangled from these data alone and will require computational modeling and multimodal integration. Fourth, source-imaging resolution and spatial leakage can influence local spatial gradients; although our surface-based formulation, artifact-control analyses, and cross-dataset replication mitigate this concern, future work should quantify sensitivity to inverse-model choice and spatial smoothing more explicitly.

## Conclusion

Geodesic cortical flow provides a non-invasive, geometry-aware framework for quantifying the direction, strength, and temporal stability of spontaneous cortical propagation at millisecond timescales. In resting-state MEG, this framework reveals frequency-specific propagation aligned with cortical hierarchy and systematically reweighted across adulthood. These measures may offer a useful route toward in vivo markers of cortical organization, aging, and neurocognitive vulnerability.

## Materials and Methods

### Participants and datasets

#### Cambridge Centre for Aging and Neuroscience (Cam-CAN)

To characterize how spontaneous cortical propagation varies across the adult lifespan, we analyzed resting-state data from the Cambridge Centre for Ageing and Neuroscience (Cam-CAN) cohort (Taylor et al., 2017). The sample comprised 608 healthy adults aged 18–88 years (307 men, 301 women; mean age 54.19 ± 18.19 years), each with resting-state MEG and T1-weighted MRI.

MEG was recorded during eyes-closed rest on a whole-head MEGIN system (Helsinki, Finland) at 1000 Hz. Cognitive performance was indexed by the Cam-CAN fluid-intelligence composite (mean ± SD, 31.80 ± 6.79), derived from four subtests: series completion, classification, matrices, and conditions. This cohort allowed us to test age-related variation in cortical-flow directionality, kinetic properties, and cognitive associations.

#### Open MEG Archive (OMEGA)

To assess robustness across acquisition systems, we analyzed an independent resting-state dataset from the Open MEG Archive (OMEGA; Niso et al., 2016). After quality control and completeness checks, the final sample included 83 healthy adults (40 women, 43 men; mean age 28.66 ± 6.97 years). MEG was acquired with a 275-channel whole-head CTF system (Port Coquitlam, BC, Canada) at 2400 Hz. Unless noted otherwise, the same preprocessing and analysis pipeline was applied as in Cam-CAN.

### MEG and MRI preprocessing

#### MEG preprocessing

All preprocessing was performed in *Brainstorm* (Tadel et al., 2011) using default settings unless otherwise specified and in accordance with current MEG reporting guidelines (Gross et al., 2013).

MEG data were band-pass filtered between 0.3 and 200 Hz with finite impulse response filters to remove slow drift and high-frequency noise. Line noise and its harmonics were removed with notch filters (OMEGA: 60, 120, and 180 Hz; Cam-CAN: 50, 100, and 150 Hz). Bad channels and noisy segments were identified manually and excluded. Ocular and cardiac artifacts were attenuated with signal-space projection (SSP) using projectors derived from EOG- and ECG-locked epochs.

Power spectral density (PSD) was estimated for each participant with Welch’s method using 2-s windows and 50% overlap. To define an adaptive upper frequency limit for subsequent analyses, we identified the frequency below which 95% of total PSD power was contained. Data were then segmented into 30-s non-overlapping epochs to ensure stable estimation of resting-state dynamics (da Silva Castanheira et al., 2021; Wiesman et al., 2022). Each epoch was resampled at four times the participant-specific PSD cut-off to reduce memory demand while avoiding aliasing (Michel & Brunet, 2019).

#### MRI preprocessing and MEG source reconstruction

Structural MRI preprocessing was performed with *FreeSurfer* (Dale et al., 1999) to reconstruct the cortical surfaces and generate triangular meshes. MEG and MRI data were coregistered using approximately 100 digitized scalp points. Forward models were computed with the overlapping-spheres method, and source time series were reconstructed with linearly constrained minimum variance beamforming with depth-bias correction. Noise covariance was estimated from empty-room recordings.

Source time series were extracted for broadband activity (up to the participant-specific PSD cut-off) and for canonical frequency bands. Primary analyses focused on slow activity (1–13 Hz) and beta activity (13–30 Hz), using zero-phase finite impulse response filtering. Boundary robustness was checked by shifting the band boundary by ±1 Hz.

### Cortical-flow estimation on the cortical manifold

We modeled the cortical surface as a two-dimensional Riemannian manifold embedded in three-dimensional space (Supplementary Fig. 2) and discretized as a triangular mesh. Cortical activity was treated as a scalar field evolving over time on this manifold.

In local coordinates, the surface normal at each vertex is defined by the cross-product of the partial derivatives of the embedding function:

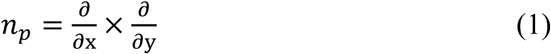

Because this normal depends only on local geometry, it is invariant to the choice of coordinate system used for parameterization.

Let I denote the scalar field representing cortical activity (for example, the source-imaged MEG signal). Its differential maps tangent vectors on the cortical manifold to real values and provides the local directional variation needed to define spatial gradients and cortical flow.

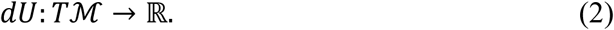

For two tangent vectors, we define their differential interaction as follows:

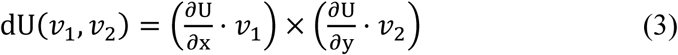

This formulation provides the geometric ingredients used to estimate local propagation directly on the cortical surface.

Cortical flow is conceptually analogous to optical flow in computer vision, where apparent motion is inferred from successive image frames (Lefèvre et al., 2008). Here, cortical activity is treated as a sequence of scalar maps defined on the cortical surface. Assuming local conservation of this scalar field along flow trajectories, the cortical flow field satisfies the transport equation:

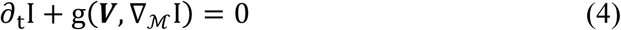

Here, the Riemannian metric g(.,.) modifies the local inner product according to surface curvature (Lefèvre et al., 2008). This conservation assumption is appropriate at MEG timescales, where activity patterns change smoothly across consecutive samples. As in classical optical-flow estimation, the transport equation alone does not uniquely determine the flow field in regions where gradients are locally ambiguous (the aperture problem). We therefore estimated the cortical-flow vector field by minimizing an energy functional that balances fidelity to the observed dynamics with spatial smoothness:

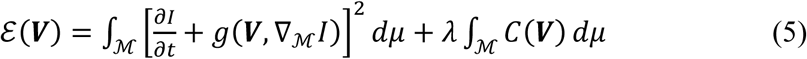

In this expression, the manifold volume form determines local integration weights, λ controls the trade-off between data fidelity and smoothness as a regularization term penalizing spatially abrupt changes in the estimated vector field. We set λ = 0.01, following Lefèvre et al. (2008).

### Anatomy-informed geodesic reference frame

To enable consistent interpretation of propagation direction across the folded cortical surface, we defined an anatomy-informed geodesic reference frame (Fig. 1B; Supplementary Fig. 2). Individual cortical surfaces and flow fields were registered to the *fsaverage* spherical template, which standardized vertex correspondence while preserving continuous surface trajectories.

At each vertex, a local Cartesian coordinate system was defined in the tangent plane. A sagittal anterior-posterior axis was constructed separately for each hemisphere using the most anterior and most posterior cortical vertices in template space; an orthogonal superior-inferior axis was then defined within the tangent plane. Hemisphere-specific adjustments ensured consistent anatomical orientation across medial and lateral surfaces. Flow vectors were projected into this frame, and propagation direction was quantified as angular deviation from canonical anatomical axes.

### Directional analyses and circular statistics

All directional analyses were performed with the *Circular Statistics Toolbox* for MATLAB (Berens, 2009). Flow vectors were expressed as angles in the local geodesic reference frame, with 90° denoting posterior-to-anterior propagation and 270° denoting anterior-to-posterior propagation. Participant-level circular means and directional distributions were then computed across time.

### Alignment with macroscale functional hierarchy

We tested whether cortical-flow vectors aligned with the principal functional gradient spanning unimodal sensory-motor regions to transmodal association cortex (Margulies et al., 2016). A normative map of this gradient was obtained with *BrainSpace* (Vos de Wael et al., 2020) and projected onto the *fsaverage* surface.

Local hierarchy direction was defined as the surface gradient of the functional-hierarchy map, yielding a unit vector field. At each vertex and time point, we computed the unsigned angular difference between cortical-flow direction and hierarchy direction on the interval [0°, 180°]. Continuous alignment was quantified with cosine similarity. For descriptive summaries, propagation was classified as hierarchy-aligned when 0° ≤ Δθ < 90° and hierarchy-opposed when 90° ≤ Δθ ≤ 180°. Metrics were summarized across vertices and time points at the participant level.

Deviation from circular uniformity was assessed with the Hodges-Ajne test, and bimodality of angular differences was assessed with Hartigan’s dip test. Statistical significance was determined with spin-permutation nulls that preserved spatial autocorrelation (1,000 rotations).

### Frequency-dependent propagation across the lifespan

To examine frequency-dependent propagation, we computed for each participant and frequency band the proportion of time points during which cortical flow was aligned with the functional hierarchy. Differences between slow and beta activity were assessed with paired-sample t tests, and effect sizes were summarized with Cohen’s d.

Age-related modulation of these propagation measures was assessed with partial correlations controlling for sex and handedness. Multiple comparisons were controlled with false discovery rate (FDR) correction.

### Kinetic energy and state-based temporal analysis

Propagation strength was quantified as kinetic energy, defined as the squared norm of the surface-tangent flow vector at each vertex and time point. Global kinetic energy was computed as the mean across vertices.

Temporal fluctuations in kinetic energy were used to separate stable states from transition states (Ashourvan et al., 2017; Watanabe et al., 2014). Stable states were centered on local minima and transition states on local maxima after band-specific smoothing, with smoothing windows derived from the relevant filter cut-offs (Leonardi et al., 2015; Preti et al., 2017). Stable-state dwell time was defined as the duration of contiguous low-energy epochs.

### Propagation kinetics and cortical temporal hierarchies

We assessed whether propagation kinetics tracked known cortical hierarchies of functional specialization and temporal organization. Spatial correspondence was evaluated relative to the principal functional gradient (Margulies et al., 2016) and to a T1w/T2w-derived myelination index (Glasser & Van Essen, 2011), with significance assessed by spin permutation.

To relate propagation stability to intrinsic neuronal timescales, we estimated spectral knee parameters with *specparam* (Donoghue et al., 2020). Intrinsic timescales were derived from the knee frequencies and compared with stable-state dwell times across cortical regions defined following Murray et al. (2014) and Gao et al. (2020). Vertex-wise maps of dwell time and intrinsic timescale were also spatially correlated and tested with spin permutations.

### Frequency-specific kinetics of cortical propagation

Beyond broadband analyses, we examined frequency-specific differences in propagation direction and kinetics. Specifically, we tested whether slow activity (1–13 Hz) was preferentially aligned with the sensory-to-association axis and whether beta activity (13–30 Hz) was preferentially aligned with the opposite direction. Frequency-specific propagation strength was summarized with kinetic energy and with the beta-to-slow kinetic-energy ratio.

### Age and cognition analyses

Associations between kinetic-energy measures and age were tested with correlation and regression models that included sex, handedness, head motion, and residual physiological components as covariates. Relationships with fluid intelligence were assessed using age-adjusted residuals. Multiple comparisons were controlled with FDR correction, and spatial correspondence was evaluated with spin permutations (Alexander-Bloch et al., 2018; Markello & Misic, 2021) based on 1,000 rotations.

## Data availability

The raw MEG and structural MRI data used in this study are publicly available from the Cambridge Centre for Ageing and Neuroscience (Cam-CAN; https://www.cam-can.org/) and the Open MEG Archive (OMEGA; https://www.mcgill.ca/bic/neuroinformatics/omega). Resting-state fMRI data used to derive the normative functional gradient were obtained from the Human Connectome Project (HCP; https://www.humanconnectome.org/).

The gradient map of the cortical functional hierarchy was generated with the BrainSpace toolbox: https://brainspace.readthedocs.io/en/latest/.

Supplementary movies illustrating the dynamic features described in this study are available at: https://drive.google.com/drive/folders/1s_reaRJlw9uyJvNGG39M7RTIJsAPgCxL.

## Code availability

Analysis code is available at https://github.com/Laoma29/Cortical-Flow-Project.

## Acknowledgments

Xiaobo Liu was supported by the China Scholarship Council. Alex I. Wiesman is supported by the Tier-2 Canada Research Chair in Neurophysiology of Aging and Neurodegeneration (CRC-2023-00300). Sylvain Baillet was supported by a Discovery Grant from the Natural Sciences and Engineering Research Council of Canada (436355-13), the CIHR Canada Research Chair in Neural Dynamics of Brain Systems (CRC-2017-00311), the NIH (R01-EB026299-05), and the Canadian Institutes of Health Research (202503PJT-542788).

## Supplementary Material

### Validation datasets: simulations, artifact controls, and replication

To validate the cortical-flow framework, we combined synthetic simulations with known ground-truth propagation, targeted artifact-control analyses, and replication of the core findings in an independent dataset (OMEGA).

### Methodological validation: simulation of cortical propagation patterns

To evaluate directional accuracy, we simulated biologically plausible cortical propagation on a curved surface using the vectorial heat equation (Lefèvre et al., 2009). This framework extends the classical scalar heat equation to vector fields defined on manifolds and provides smooth, spatially coherent trajectories across cortical geometry.

We begin with the classical parabolic partial differential equation (PDE) defined on a 2D Euclidean domain Ω:

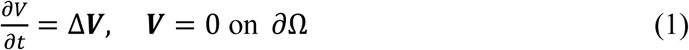

Where the vectorial Laplacian is defined as:

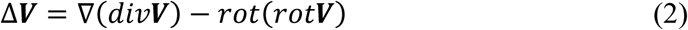

Rather than computing this expression explicitly on the manifold *M*, we adopt a variational formulation of the vectorial heat equation, which is more computationally tractable and enables numerical integration over curved surfaces. The evolution of the vector field ***V*** is governed by:

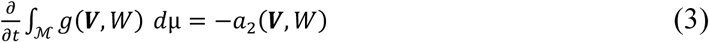

where *g*(⋅,⋅) denotes the Riemannian metric, *d*μ is the volume form on *M* , *W* ∈ *T*ℳ is a test vector field, and *a*_2_(***V***, *W*) is a bilinear form involving covariant derivatives of ***V*** and *W*. This vectorial diffusion process promotes smoothness and spatial coherence, properties required for biologically realistic simulations of cortical activity propagation.

Using this framework, we simulated large-scale propagation patterns along three cardinal axes: posterior→anterior, anterior→posterior, and inferior→superior (Figure 1). Ground-truth direction at each vertex was defined by the displacement vector between successive time points. To assess statistical significance, we implemented a hemisphere-constrained spherical rotation (“spin”) null model that preserves spatial autocorrelation. Each trajectory was randomly rotated 1,000 times across the cortical surface, generating a null distribution of directional vectors. Observed mean directions and angular deviations were compared against this null, yielding empirical p_spin_ values.

Cortical-flow estimates recovered the ground-truth directions across all simulated conditions (Supplementary Fig. 3). With 90° denoting posterior-to-anterior and 0° denoting superior-to-inferior directions, the posterior-to-anterior simulation yielded a mean estimated direction of 86.81° (95% CI, 73.64°–99.99°; mean angular error 0.97°), the anterior-to-posterior simulation yielded 275.69° (95% CI, 259.51°–291.85°; mean angular error 3.67°), and the inferior-to-superior simulation yielded 157.56° (95% CI, 142.09°–173.61°; mean angular error 11.81°). All recovered directions differed significantly from the spin-based null (pspin < 0.05).

### Artifact control analysis

To evaluate whether transient physiological artifacts influenced cortical-flow direction or kinetic energy, we performed targeted control analyses using epochs locked to eyeblinks and heartbeats in the post-QC OMEGA sample (N = 83). Blink onsets were identified from the vertical EOG, and heartbeat onsets were identified from ECG.

For each blink, we extracted epochs spanning -300 to +300 ms and processed the resulting eye-movement event-related fields with the cortical-flow framework. Global and parcellated flow directions were compared between these epochs and artifact-free data, and local directional structure was assessed with the Hodges-Ajne test using FDR correction across regions.

We repeated the same procedure for cardiac artifacts using epochs spanning -50 to +50 ms around each R-peak.

To assess preprocessing effects directly, cortical-flow metrics were also compared before and after signal-space projection (SSP) within artifact-free resting segments.

Blink epochs exhibited a reproducible biphasic pattern, with anterior-to-posterior propagation at blink onset followed by a posterior-to-anterior rebound at blink offset (Supplementary Movie 8). Heartbeat epochs showed the converse pattern (Supplementary Movie 9). Mean onset angles were 15.1° (95% CI, 13.6°–16.5°) for blinks and 188.7° (95% CI, 186.0°–191.3°) for heartbeats.

After SSP correction, this directional structure was abolished and angular distributions no longer differed from uniformity (pperm > 0.05). Kinetic-energy transients visible around blink and heartbeat events were likewise removed, and post-SSP values no longer differed from artifact-free resting segments (paired t test, p > 0.05).

These analyses indicate that physiological artifacts can induce spurious propagation and kinetic-energy transients, but that SSP effectively removes these effects.

### Replication analysis (OMEGA)

To test generalizability, we repeated the core analyses in an independent resting-state MEG sample from the OMEGA repository (N = 83), using identical preprocessing and metric definitions.

Broadband activity (0.6–92.6 Hz) again exhibited sagittal posterior-to-anterior propagation, with a group-mean angle of 74.6° (95% CI: 72.8°–76.4°, p_perm_ < 0.001). Flow vectors remained preferentially aligned with the principal functional hierarchy, and angular differences showed a bimodal distribution (Hartigan dip, p_spin_ < 0.001), indicating co-existing hierarchy-aligned and hierarchy-opposed streams.

Frequency-specific bias replicated: slow activity (1–13 Hz) showed a higher incidence of hierarchy-aligned propagation than beta activity (13–30 Hz) (paired *t* = 2.80, *p* = 0.006, *Cohen’s d* = 0.35).

The spatial pattern of kinetic energy mirrored Cam-CAN, with higher values in posterior sensory cortex decreasing toward anterior association areas, yielding a negative correlation with functional hierarchy rank (*r* = −0.66, *p_spin_* < 0.001). Kinetic energy also correlated negatively with a myelination index derived from T1w/T2w ratios (*r* = −0.66, *p_spin_* < 0.001).

Stable-state dwell times increased along the same posterior-to-anterior hierarchy (*r* = 0.30, *p_spin_* < 0.001) and correlated positively with intrinsic neuronal timescales extracted from spectral knees (*r* = 0.51, *p_spin_* < 0.001).

Together, the OMEGA replication confirms that sagittal propagation, hierarchy alignment, frequency-specific directional biases, and frequency-resolved kinetic gradients generalize across datasets and acquisition platforms.

---

**Supplementary Figure 1:**
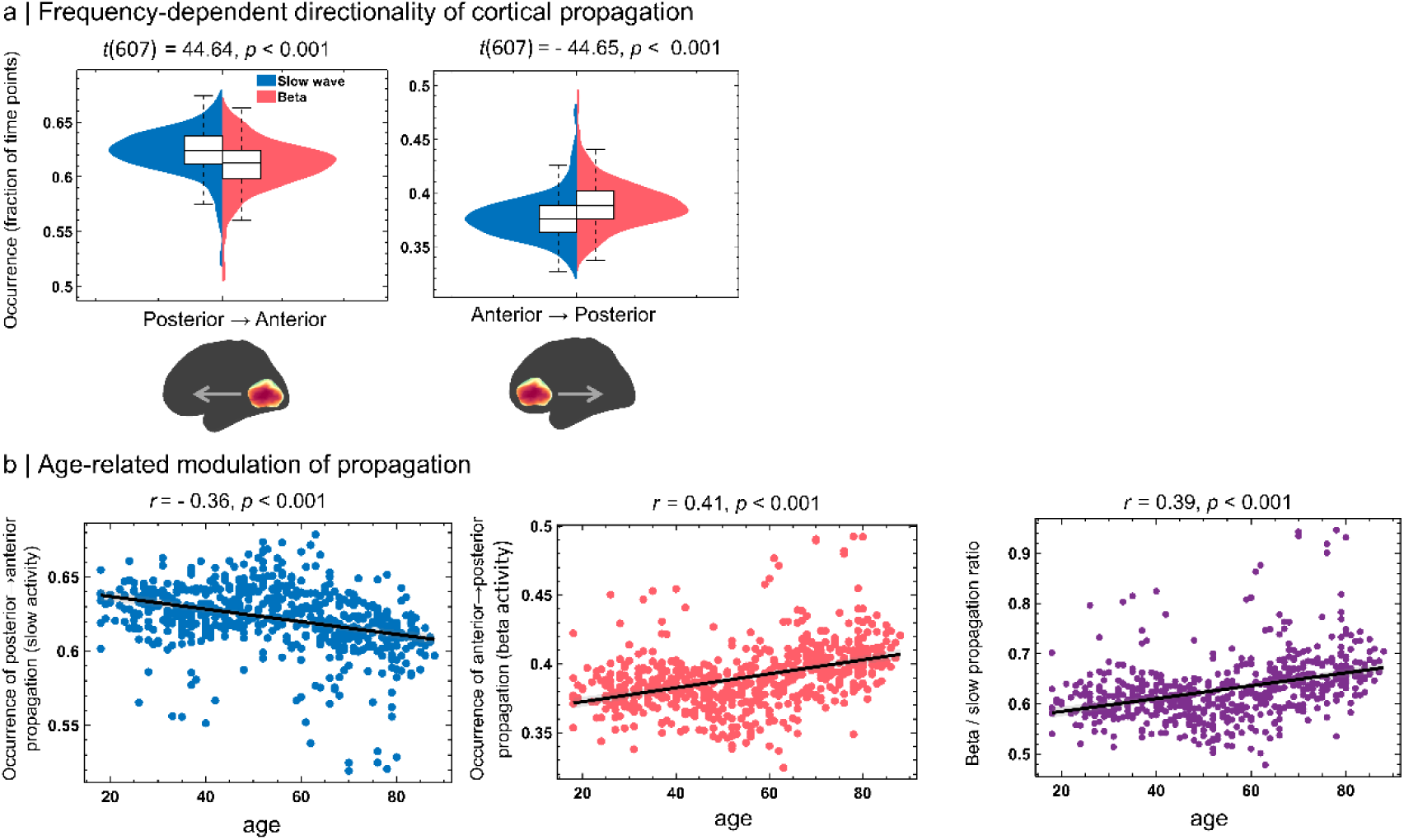
Age-related changes in cortical propagation direction across frequency bands. **a | Conceptual model of frequency-specific propagation patterns:** This schematic illustrates the hypothesized dominant propagation trajectories of spontaneous cortical activity across frequency bands. Slow-frequency activity (blue) predominantly propagates from posterior to anterior regions, reflecting bottom-up integration, while β-frequency activity (red) tends to propagate in the opposite direction, consistent with top-down control. **b | Frequency-specific propagation direction distributions:** Violin plots show the group-level distributions of propagation direction frequencies across participants, comparing slow and β bands along two sagittal axes: • Left: Slow activity exhibits significantly more frequent posterior-to-anterior propagation (*t* = 44.64, *p* < 0.0001). • Right: Β activity shows significantly greater anterior-to-posterior propagation (*t* = –44.65, *p* < 0.0001). These findings confirm a frequency-dependent reversal in preferred propagation direction. **c | Lifespan changes in propagation direction:** Scatter plots display how propagation direction varies with age: • Left: Posterior-to-anterior propagation of slow activity declines with age (*r* = – 0.36, *p* < 0.0001). • Middle: Anterior-to-posterior propagation of β activity increases with age (*r* = 0.41, *p* < 0.0001). • Right: The β-to-slow directional ratio increases with age (*r* = 0.39, *p* < 0.0001), indicating a shift toward top-down dynamics in aging.

**Supplementary Figure 2:**
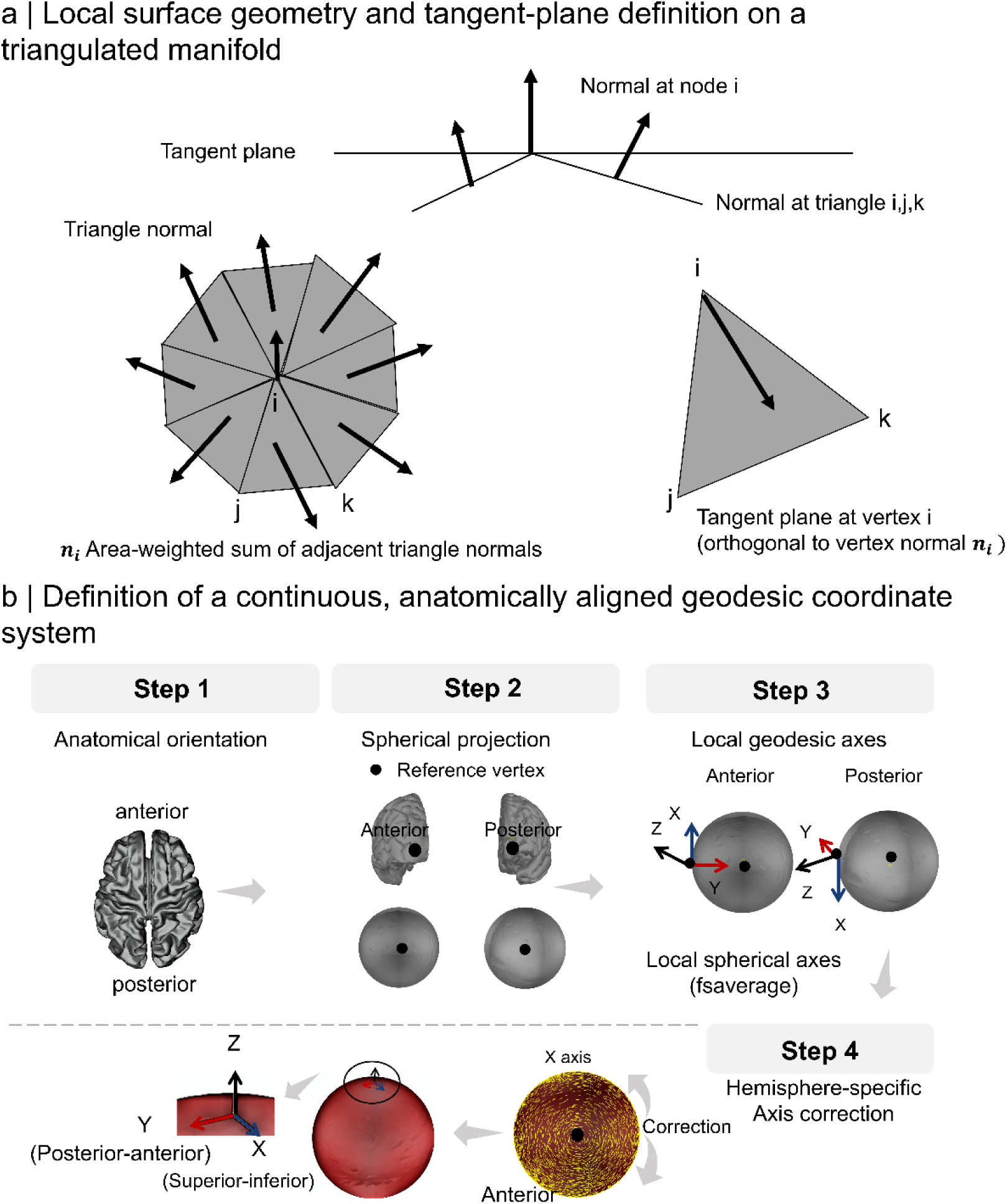
Cortical Riemannian manifold and geodesic coordinate system. **a | Riemannian manifold:** Schematic illustration of a 2D Riemannian manifold discretized as a triangular cortical mesh in 3D space. At each vertex, a local tangent plane is computed from neighboring faces. The surface normal vector n_p_ is derived via the cross product of local basis vectors in the plane. This construction enables the application of differential operators, such as gradients and vector field flow, to characterize propagation of neural activity across the surface. **b** | **Geodesic coordinate system:** This figure illustrates the procedure used to define a consistent, anatomically grounded reference frame for computing and interpreting cortical flow vectors across the cortical surface. For each hemisphere, we defined the posterior-anterior axis using the most posterior and most anterior vertices on the *fsaverage* surface. These two points define a posterior-to-anterior Y-axis, which is then projected onto the spherical representation of the cortical manifold. Next, a surface-based X-axis is defined as orthogonal to the Y-axis in the superior–inferior direction, and a Z-axis is derived accordingly to complete the local 3D Cartesian coordinate system (X, Y, Z) at each vertex. To preserve directional consistency across hemispheres—particularly across lateral and medial surfaces—a coordinate flip is applied on one hemisphere to align the anatomical orientations of X across the entire cortex. This anatomically informed, geodesic-based coordinate system enables reliable and uniform interpretation of cortical flow directionality, facilitating group-level comparisons and robust alignment with macroscale cortical hierarchies.

**Supplementary Figure 3:**
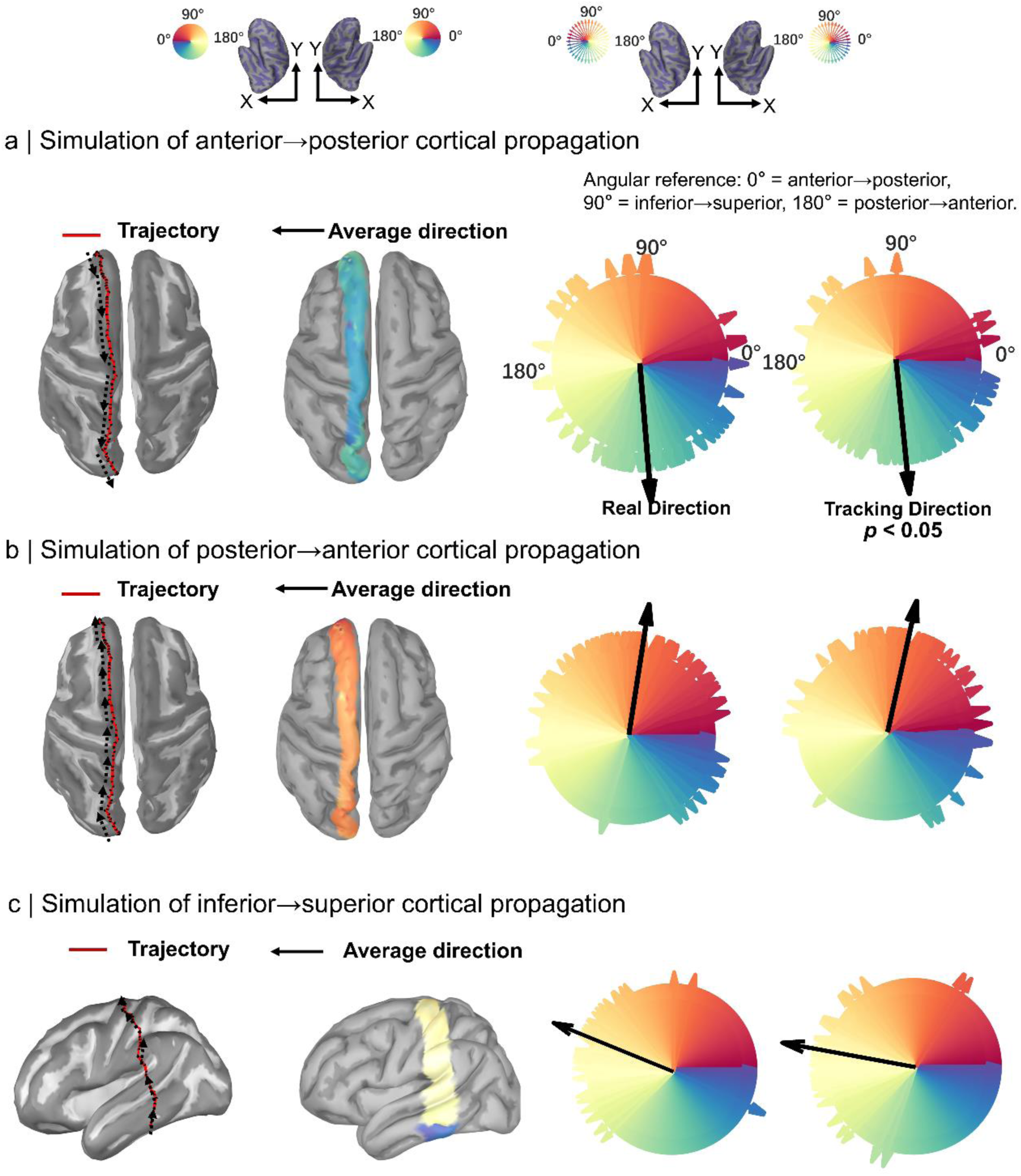
Validation of cortical-flow tracking using synthetic propagation patterns. This figure illustrates validation of the cortical-flow framework using synthetic simulations in which propagation trajectories and directions are known a priori. Synthetic activity patterns were generated by solving the vector heat equation on a Riemannian cortical manifold, producing controlled propagation along predefined anatomical axes. Each simulation therefore provides a ground-truth direction against which estimated propagation can be quantitatively assessed. **a | Anterior→posterior propagation:** Activity was initiated in frontal cortex and propagated toward the occipital pole. The estimated mean propagation angle was 275.69° (95% CI: [259.51°, 291.85°]), with a mean angular error of 3.67°. The rose plot shows strong alignment between tracked and ground-truth directions (Rayleigh test, *p* < 0.05). **b | Posterior→anterior propagation:** Activity propagated from occipital to frontal cortices. The estimated mean angle was 86.81° (95% CI: [73.64°, 99.99°]), with a mean angular error of 0.97°, indicating highly accurate directional tracking. **c | Inferior→superior propagation:** Activity propagated from ventral temporal toward dorsal parietal cortex, traversing curved, vertically oriented cortical pathways. The estimated mean angle was 157.56° (95% CI: [142.09°, 173.61°]), with a mean angular error of 11.81°, reflecting increased geometric complexity while preserving correct directional alignment.

**Supplementary Figure 4:**
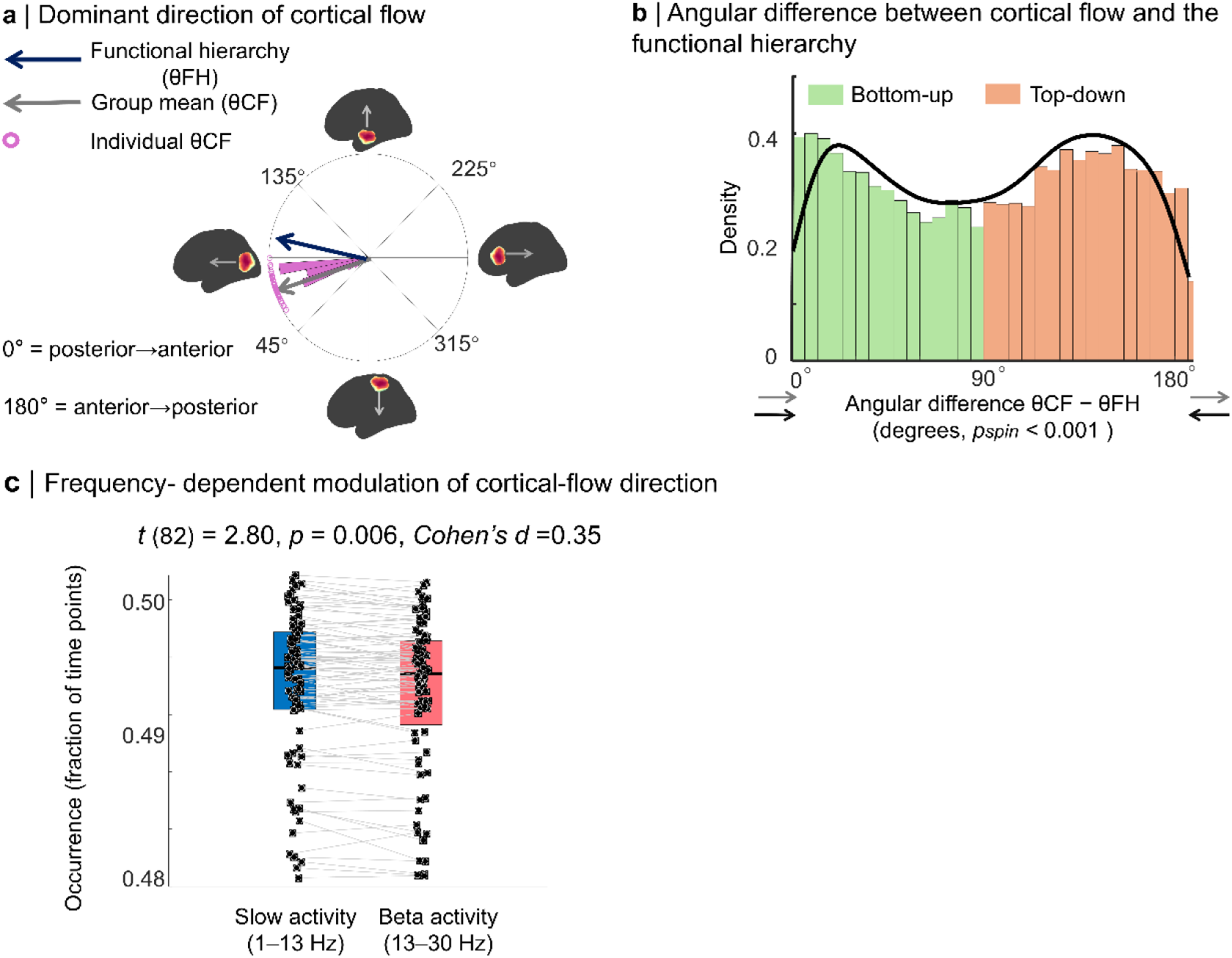
Replication of hierarchy-aligned cortical propagation in the OMEGA dataset (N = 83). **a | Dominant direction of cortical flow (**θ_CF_**) across OMEGA participants**: Polar plot shows individual participant mean cortical-flow directions (gray arrows) relative to the principal functional hierarchy direction (θ_FH_; blue arrow), revealing a predominant posterior→anterior orientation of spontaneous cortical propagation at the group level. **b | Angular difference between cortical-flow direction and the functional hierarchy:** Histogram of θ_CF_ − θ_FH_ reveals a robust bimodal distribution (*p_spin_* < 0.001), with one mode near 0° corresponding to propagation aligned with the functional hierarchy (bottom-up) and a second mode near 180° corresponding to propagation opposed to the hierarchy (top-down). c | Frequency-dependent modulation of hierarchy-aligned propagation: Paired dot-and-box plots compare the occurrence of hierarchy-aligned propagation for slow activity (1–13 Hz; blue) and beta activity (13–30 Hz; red). Slow activity shows significantly greater hierarchy-aligned propagation than beta activity (t(82) = 2.80, p = 0.006, Cohen’s d = 0.35), replicating the Cam-CAN result.

**Supplementary Figure 5:**
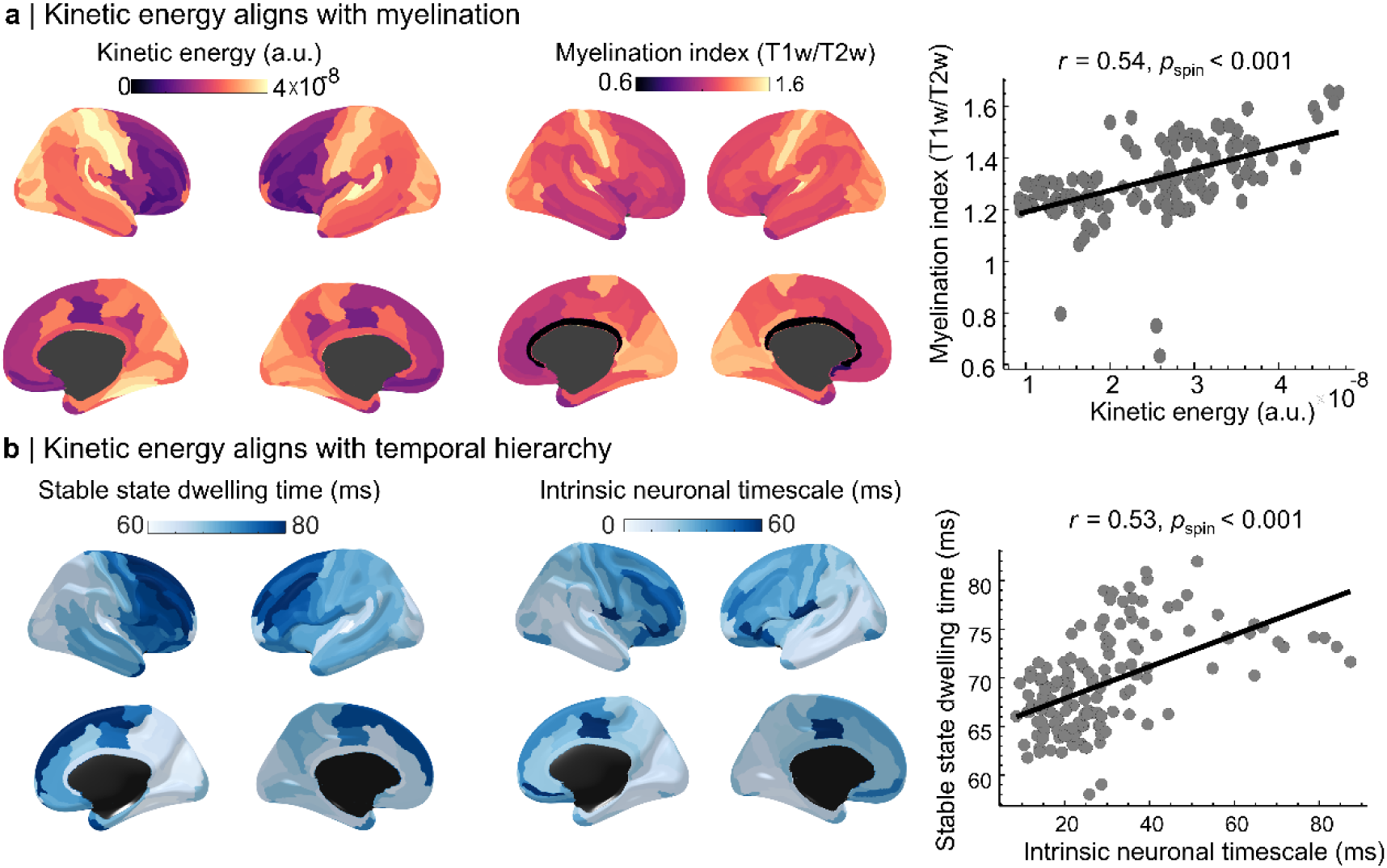
Alignment of cortical kinetic energy with myelination and neuronal timescales. **a | Relationship between cortical kinetic energy and myelination:** Surface maps show the spatial distribution of kinetic energy and a myelination index derived from T1w/T2w ratios. Vertex-wise analysis reveals a significant positive association between kinetic energy and cortical myelination (*r* = 0.54, *p_spin_* < 0.001), linking propagation strength to underlying microstructural gradients. b | Relationship between stable-state dwell time and intrinsic neuronal timescales: Surface maps show stable-state dwell time and intrinsic neuronal timescale. Stable-state dwell time correlates positively with regional neuronal timescales (r = 0.53, pspin < 0.001), supporting its alignment with the cortical temporal hierarchy.

**Supplementary Figure 6:**
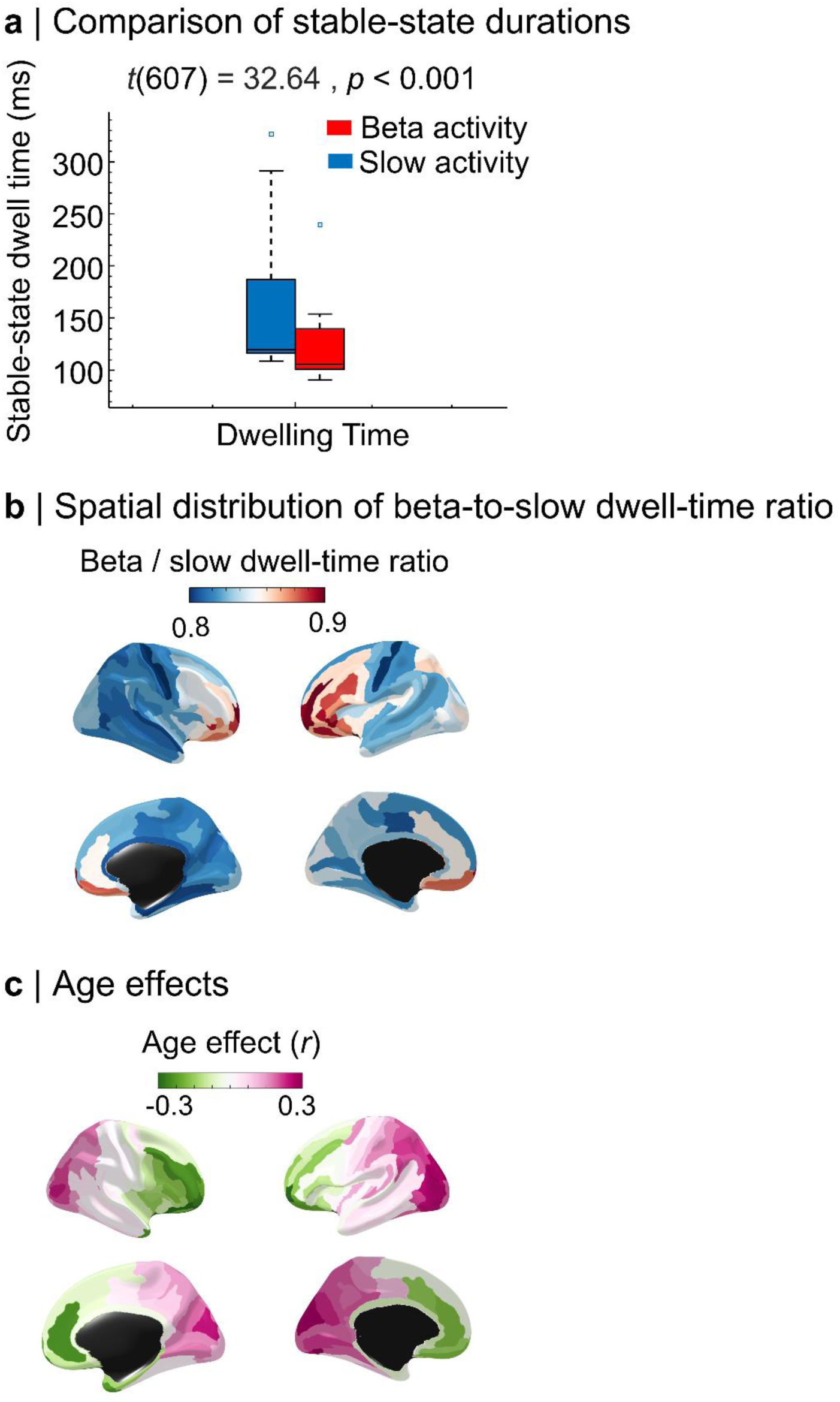
Frequency-specific differences in stable-state dwell times and their modulation with aging (Cam-CAN; N = 608). **a | Comparison of stable-state dwell times for slow (1–13 Hz) and beta-band (13–30 Hz) activity:** Stable states persist significantly longer for slow activity than for beta activity (*t*(607) = 32.64, *p* < 0.001), indicating that slower activity supports more sustained cortical states. **b | Spatial distribution of the beta-to-slow dwell-time ratio**: Cortical maps show the ratio of beta-band to slow-activity stable-state dwell times across the cortex. Higher ratios in association cortex indicate relatively greater stability of beta-band states compared to slow activity, revealing a spatial organization of frequency-specific stability. **c | Age-related modulation of the beta-to-slow dwell-time ratio**: Vertex-wise analysis reveals significant associations between age and the beta-to-slow dwell-time ratio (*p_FDR_* < 0.05), indicating systematic shifts in the balance between fast and slow stable-state dynamics across the adult lifespan.

